# Massive Factorial Design Untangles Coding Sequences Determinants of Translation Efficacy

**DOI:** 10.1101/208801

**Authors:** Guillaume Cambray, Joao C. Guimaraes, Adam Paul Arkin

**Affiliations:** California Institute for Quantitative Biosciences, University of California, Berkeley, CA,94720, USA.; DGIMI, INRA, Univ. Montpellier, Montpellier, France.; Department of Bioengineering, University of California, Berkeley, CA, 94720, USA.; Physical Biosciences Division, Lawrence Berkeley National Laboratory, Berkeley, CA, 94720, USA.

## Abstract

Comparative analyses of natural sequences or variant libraries are often used to infer mechanisms of expression, activity and evolution. Contingent selective histories and small sample sizes can profoundly bias such approaches. Both limitations can be lifted using precise design of large-scale DNA synthesis. Here, we precisely design 5 *E. coli* genomes worth of synthetic DNA to untangle the relative contributions of 8 interlaced sequence properties described independently as major determinants of translation in *Escherichia coli*. To expose hierarchical effects, we engineer an inducible translational coupling device enabling epigenetic disruption of mRNA secondary structures. We find that properties commonly believed to modulate translation generally explain less than a third of the variation in protein production. We describe dominant effects of mRNA structures over codon composition on both initiation and elongation, and previously uncharacterized relationships among factors controlling translation. These results advance our understanding of translation efficiency and expose critical design challenges.

## INTRODUCTION

The modulation of translation rate is a major contributor to global and individual gene regulation programs (Vogel and Marcotte, 2012). In itself, the translational machinery represents *ca*. 40% of the total protein biosynthesis (Li et al., 2014), so that its pattern of utilization is intricately linked to the physiological state of the cell (Scott et al., 2010). Although the evolution of coding sequences is primarily constrained by the necessity to maintain protein biochemical functions, it has become clear that functional redundancies—notably within the genetic code—provide space for encoding additional layers of regulatory information (Plotkin and Kudla, 2010). A better understanding of the sequence properties capable of impacting translation rates has therefore important fundamental and biotechnological implications. Nonetheless, their exact nature and respective prevalence have been subject to much debate (Gingold and Pilpel, 2011; Tuller and Zur, 2015).

The early accumulation of gene sequences swiftly revealed species-specific biases in the genomic usage of synonymous codons (Grantham et al., 1980). The realization that the codon composition of the most highly expressed genes in a given genome correlate tightly with the cellular abundance of cognate tRNAs (Ikemura, 1985) stirred the development of metrics aimed at predicting the expression level of genes from their sequences alone (Cannarozzi and Schneider, 2012). For example, the widely used Codon Adaptation Index (CAI, Sharp and Li, 1987) provides as good a predictor of protein abundances as experimentally measured mRNA in *E. coli* (r=0.54 and 0.56, respectively, using n=575 from Taniguchi et al., 2010). As a result, this and related metrics became *de facto* gold standards for genomic prediction of expression and—by extension—biotechnological applications (Gustafsson et al., 2004).

With the scaling of genetic engineering capacities, a number of experimental results challenged the capacity of such metrics to support rational design of heterologous gene expression. Instead, numerous mechanistic studies have highlighted the major role of mRNA secondary structures in occluding important sequence signals from the ribosomes and consequently decrease translation initiation rates (Adhin and van Duin, 1990; Espah Borujeni et al., 2014; Kudla et al., 2009; Mutalik et al., 2013a). Controlling for such structures by design, Allert et al. (2010) assayed 285 synthetic genes *in vitro* and concluded that %AT at the beginning of coding sequences had a larger impact on protein production than structures themselves. Because the distributions of these sequence properties are highly biased around the start of natural coding sequences (Gu et al., 2010), their functional importance had long been overlooked by traditional genomics correlation analyses (Supek and Šmuc, 2010; Tuller et al., 2010a).

In addition, an average ramp of rare and presumably slowly translated codons at the beginning of coding sequences has been described in many organisms (Tuller et al., 2010b). This unexpected feature was suggested to help distribute ribosomes along transcripts, and optimize protein production by avoiding ribosome stacking (Navon and Pilpel, 2011). Although genome-wide ribosome profiling data show higher ribosomal occupancies at the beginning of natural genes (Ingolia et al., 2009), they largely fail to validate a generic correspondence between codon usage and translational velocity. Instead, a variety of other sequence properties were found to correlate with these ribosome occupancies *e.g*. occurrences of Shine-Dalgarno-like sequences (Li et al., 2012), secondary structures downstream of the ribosome (Pop et al., 2014) and charge of amino acids in the ribosome’s exit tunnel (Charneski and Hurst, 2013). Again, all these properties are uniquely biased at the beginning of genes and were suggested to contribute to the translation speed ramp (Tuller et al., 2011).

Recent debates have therefore focused on unraveling the causes and consequences of compositional biases observed in the first 30-50 codons of genes. This region has been hailed as critical for fine-tuning the balance between rates of translation initiation and elongation (Eyre-Walker and Bulmer, 1993; Tuller and Zur, 2015). In the largest manipulation of codon composition to date, Goodman *et al.* designed 13 synonymous variants of the first 10 codons of 137 genes in *E. coli* and fused the recoded sequence to a standard reporter. Expression patterns amongst the ~1,800 resulting constructs confirmed that rare codons in this region are associated with increased protein production, as predicted by the codon-ramp hypothesis. However, *post-hoc* analyses of this dataset also revealed that these rare codons are AT-rich and tend to yield weaker predicted secondary structures. The apparent correlation between protein expression and codon usage would thus be confounded by the positive impact of weaker secondary structures on translation initiation (Bentele et al., 2013; Goodman et al., 2013)—or possibly by the distinct positive effect of %AT (Allert et al., 2010). Likewise, the excess of positively charged amino acids at the beginning of genes could be incidental to unrelated topological constraints related to the insertion of transmembrane proteins rather than the optimization of translational velocity (Charneski and Hurst, 2014).

**Figure 1.**
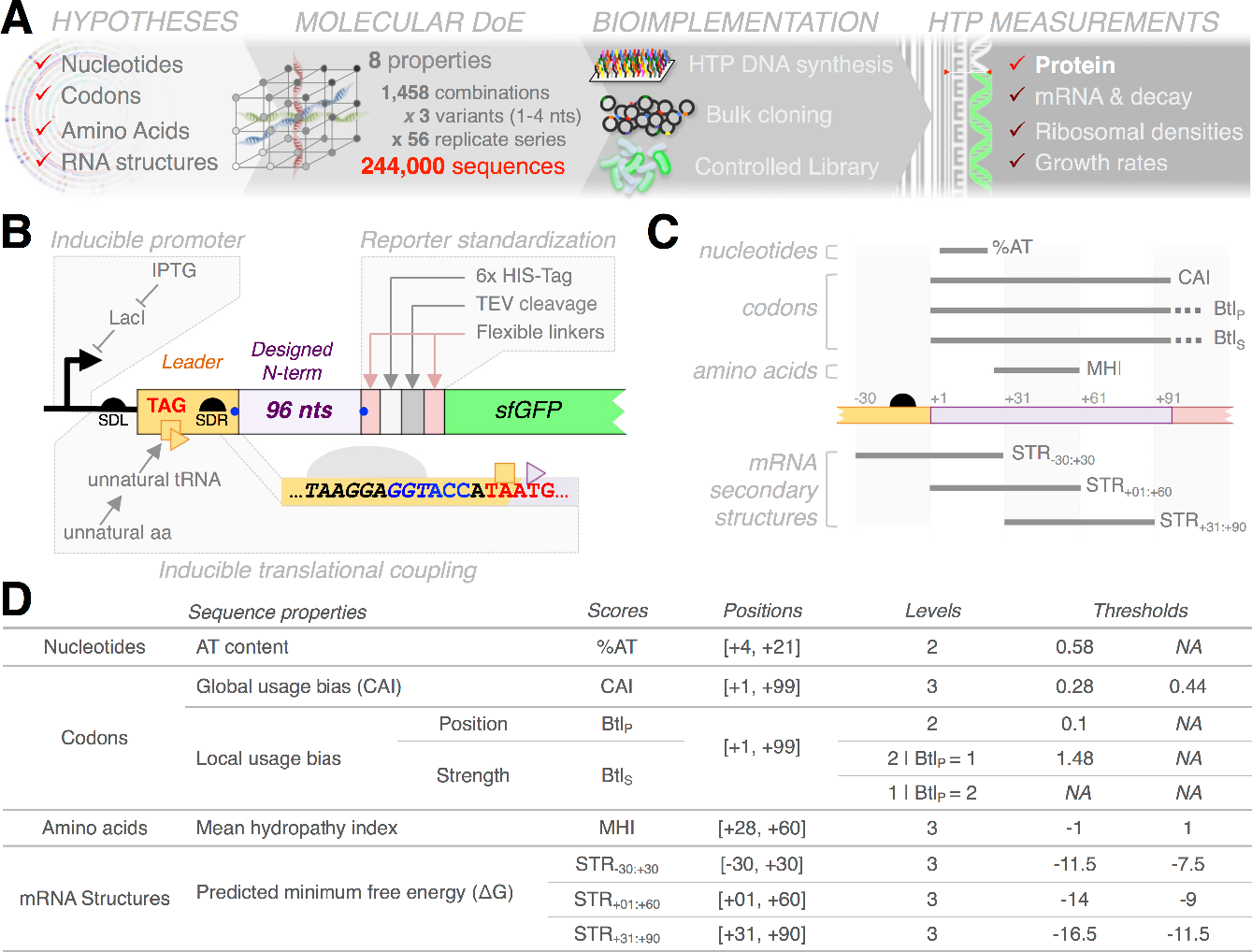
Hypothesis-driven functional genomics of translation. **(A)** High-throughput molecular Design of Experiments. Sequences were designed to produce full-factorial libraries of 8 sequence properties correlating with protein production in *E. coli*. Designer sequences were synthesized and cloned in bulk. Various phenotypes can be characterized using high-throughput sequencing as a generic quantitative readout. Here, we report on protein production. Other phenotypes are analyzed in a companion manuscript. **(B)** A standard reporter for translational output. Synthetic sequences are cloned as N-terminal fusions to a modified *sfGFP*, which produces an invariant fluorescent reporter upon post-translational processing by the TEV protease. Translation of the reporter is driven by a perfect Shine-Dalgarno motif (SDR, italic text) embedded into a novel inducible translational coupling device. Secondary structures in the initiation region can be controllably disrupted by adjusting the influx of leader-bound ribosomes through tunable induction of an unnatural suppressor system. **(CD)** Topology **(C)** and definition **(D)** of the sequence properties varied in the factorial design.

The diversity of conclusions in these reports is symptomatic of the complexity of the physical processes determining translational efficiency. They also highlight the necessity of distinguishing between arguments based on mechanistic phenomena and their evolutionary repercussions. The coding sequence properties controlling translation collectively undergo strong selective pressures to optimize efficiency on a global cellular level, especially in fast-growing microorganisms (Andersson and Kurland, 1990; Kudla et al., 2009). Although the analysis of evolved genomes permit the identification of functionally important regions and motifs, the very same patterns entangle the effect of many overlapping sequence properties on intricately related processes. This renders *ad hoc* inferences of direct causation challenging and subject to much interpretation. The manipulation of gratuitous reporters, in contrast, enables precise characterization of arbitrary sequences in a controlled genetic context. In particular, molecular Design of Experiments (mDoEs) that rationally varies combinations of interrelated sequence properties permit more definite testing of presumed multifactorial relationships (Allert et al., 2010; Guimaraes et al., 2014; Mutalik et al., 2013b; Smanski et al., 2014; Welch et al., 2009). In practice, however, such endeavors have been limited by the capacity to implement libraries that are large enough to disentangle more than a few properties, around more than a few reference sequences.

To address these limitations, we designed 244,000 synthetic sequences to systematically explore an 8-dimensional space defined from the main sequence properties discussed above. We implemented a library organized into 56 independent series of phylogenetically related sequences that instantiate as many replicates of a full factorial design. We used high-throughput sequencing to measure multiple phenotypic consequences of these sequence perturbations in a highly parallel fashion. For each strain, we quantified reporter transcript abundances and decay, polysome profiles, protein production and growth rates (Figure 1A). This rich and uniquely structured dataset reveals pervading and intricate associations between design variables and phenotypic responses, and further exposes areas in which we are missing critical understanding.

In this paper, we detail our design framework and dissect the impact of the designer sequence perturbations on protein production in different conditions of translational coupling. A more systemic analysis integrating these results with the other phenotypic responses is presented in a companion paper.

## RESULTS

### Design of experiments for hypothesis-driven functional genomics

We constructed a library of *E. coli* cells in which 8 sequence properties were systematically varied to assess their joint impact on translation efficiency. These features are encoded in the first 96 nucleotides of a fluorescent reporter gene fusion (Figure 1BC). The library conforms to a full-factorial design and therefore represents a statistically ideal sample to untangle the functional contribution of each sequence properties and their deeper interactions to protein production. To limit the influence of post-translational effects on fluorescent readout (*e.g*. differential protein folding or stability), variable N-termini are cleaved from the translated reporter *in vivo* using a site-specific protease (Kapust and Waugh, 2000). The reporter system is driven by a standard inducible promoter and is equipped with a controllable translational coupling system (figure 1B).

Sequences properties of interest were selected amongst the most used and discussed in the literature (Figure 1CD). We computed these properties for all genes of the *E. coli* MG1655 genome and examined their respective correlation with protein expression. This analysis permitted refinement of property scores and guided the definition of biologically meaningful thresholds for binning the continuous properties into categorical levels (Figure 1D, S1 and 2D). This defined a full-factorial design comprising 1,458 unique combinations. Nearly half these combinations are not represented in the natural *E. coli* genome, and randomly generated sequences also produce a very skewed sampling of this discretized property space (Figure 2AB).

**Figure 2.**
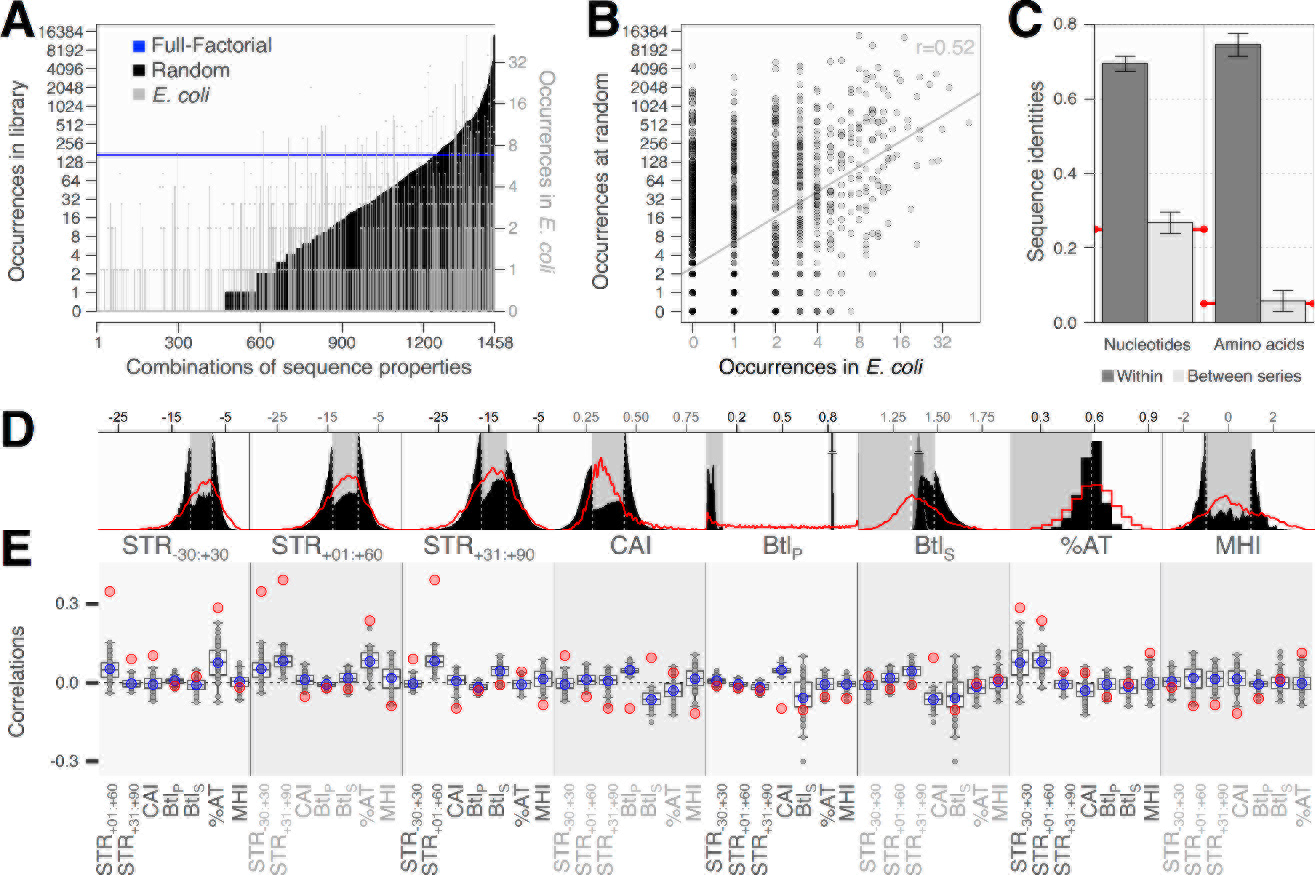
Advantages of designer over natural or random sequences. **(A)** Uneven property distributions in natural and random sequences. Black profile shows the ranked distribution of properties combinations obtained in 244,000 random sequences. Grey bars mark the corresponding distribution in natural *E. coli* genes. Both distributions are highly skewed compared to our systematic design (blue line). **(B)** Occurrences in random and natural sequences are correlated. Properties of natural sequences are partly shaped by inherent constraints and natural processes likely avoid combinations of incompatible properties. **(C)** Focal sampling of the sequence space by replicate series. The 56 full-factorial series maximize within-and minimize between-series identities at both nucleotide and amino acid levels. Shown are the mean pairwise sequences identities within (error bars show standard deviation across series) and between factorial series (error bars show standard deviation of mean identities between pairs of factorial series). Red lines mark random expectations. **(D)** Distributions of designed property scores. Designed scores are representative of wild-type *E. coli* distributions (red lines). Background shadings mark the separation between ordinal levels used for design (see table 1). Continuous scores cluster to level boundaries because extreme levels are populated from medium levels sequences during the design process. Btl_S_ nested within C-terminal Btl_P_ are shown in dark grey. **(E)** Correlations between property scores. Pairwise Pearson correlation coefficients between scores in the whole library (blue dots) or within each factorial series (grey dots and boxplots) are considerably lower than in the natural genome (red).

We thus developed a modular multi-objective sequence design framework, D-Tailor (Guimaraes et al., 2014), to generate consistent sequence libraries under such constraints. To control for factors not captured by the design parameters, we produced 56 replicate factorial series that sample distinct focal regions of the sequence space (Figure 2C and S2). To further assess the robustness of the design properties, we derived 2 close replicates for each sequence that differ by 1-4 mutations but maintain the intended combination. Details and additional constraints on design are fully described in Supplementary Information. D-Tailor was able to successfully instantiate the 244,000 sequences corresponding to this complex design. All discrete combinations of properties are equally represented in these sequences. Reflecting the difficulty to design extreme sequences, continuous property scores tend to cluster toward the design thresholds (Figure 2D). As intended, pairwise correlations between properties are drastically reduced in comparison to the reference genome (Figure 2E).

Designed sequences were synthesized on a high-density oligonucleotide array (OLS, Agilent Laboratories), PCR amplified, cloned into the reporter plasmid and transformed into *E. coli* cells (see Materials and Methods). Almost all (99.5%) of the designed sequences could be observed (≥10 reads) in a sequencing of this library targeted to the variable DNA sequences. This readily demonstrate the feasibility of implementing complex mDoE at an unprecedented scale: the combined length of sequences varied here represents more than 5 times the size of the *E. coli* genome.

### Effect of design properties on protein abundance

We used a combination of fluorescence-activated cell sorting (FACS) and targeted high-throughput sequencing to characterize fluorescent protein production as an integrated proxy for translation rate (Sharon et al., 2012; Materials and Methods). Raw data were in excellent agreement with individual measurements of more than 300 clones realized on a benchtop flow cytometer (r=0.95, Figure S3A). We could derive highly reproducible measurements of protein production (P_NC_) for 99.3% (n=242,269) of the designed sequences (median CoV=0.04 over 4 biological replicates, Figure S3BC).

We conducted a multi-way analysis of variance (ANOVA) to quantify the relative contribution of each sequence property and their second-order interactions to P_NC_ (Kosuri et al., 2013; Mutalik et al., 2013b). This procedure revealed largely dominant effect of the designed mRNA secondary structures around the start codon (STR_−30:+30_). This property alone accounts for 83% of the variability collectively explained by the design properties (Figure 3A, upper pie). A similar picture emerges from conducting a recursive regression analysis (Figure 3B).

Altogether, the design properties and their second-order interactions represent only 28% of the explainable variance, suggesting that there are important factors that are not accounted for. We therefore dissected different sources of errors in the experimental design (Figure 3A, lower pie). Variations amongst the 3 close sequence replicates implemented for each property combination within each series (median CoV=0.10, Figure S3C) account for a larger fraction of the variance than the experimental error (8% versus 1%). This design error underlines the failure of property scoring functions to capture functionally important differences between very similar nucleotide sequences (>95% identities). We also observe highly variable P_NC_ profiles amongst replicate factorial series (figure 3D). When incorporated as a categorical factor in the ANOVA, this factorial identity reclaims 10% of unexplained variance, and its primary interactions with the design properties an additional 6%. Hence, both series-specific misrepresentations of the sequence properties and unidentified sequence idiosyncrasies shared by most member of a series represent important limitations to our understanding of the system.

To better account for differences between series, we next ran independent ANOVAs on each individual factorial series (Figure 3C). The explainable variance amongst series ranges from 25% to as much as 63% (43% on average). These numbers correlate well with series-wise means and variances in P_NC_ (r=−0.42 and 0.44, respectively; Figure S3D), suggesting that poor explanation patterns may stem from inadequate implementations of key properties and consequent failure to elicit varied phenotypic responses.

Secondary structures consistently show the strongest associations with P_NC_ across all series. Specifically, STR_−30:+30_ has the largest effect size in all but two series, and accounts for 60% of explained variance on average. The structure in the very beginning of the coding sequence (STR_+01:+60_) shows the next most prevalent effect (7%), and the interaction between these two overlapping properties further contributes another 3%. Aside from structural effects, %AT and MHI account respectively for 6% and 3% of the variability on average. The remaining properties and interactions do not appear to strongly affect protein production (Figure 3B). Using continuous property scores in a multiple linear regression framework provides similar results (Figure S3EF), thereby confirming that the design thresholds do not distort the signal from the underlying properties.

Properties linked to codon usage had little impact on protein production. We could detect a slight increase in the median CAI of constructs with highest expression levels—with CAI explaining 0.75%of the variability in protein production amongst the top quintile of protein producers, versus 0.01% overall (Figure S3G). In contrast, manipulation of the putative codon ramp did not yield any measurable effect (Figure S3H).

**Figure 3.**
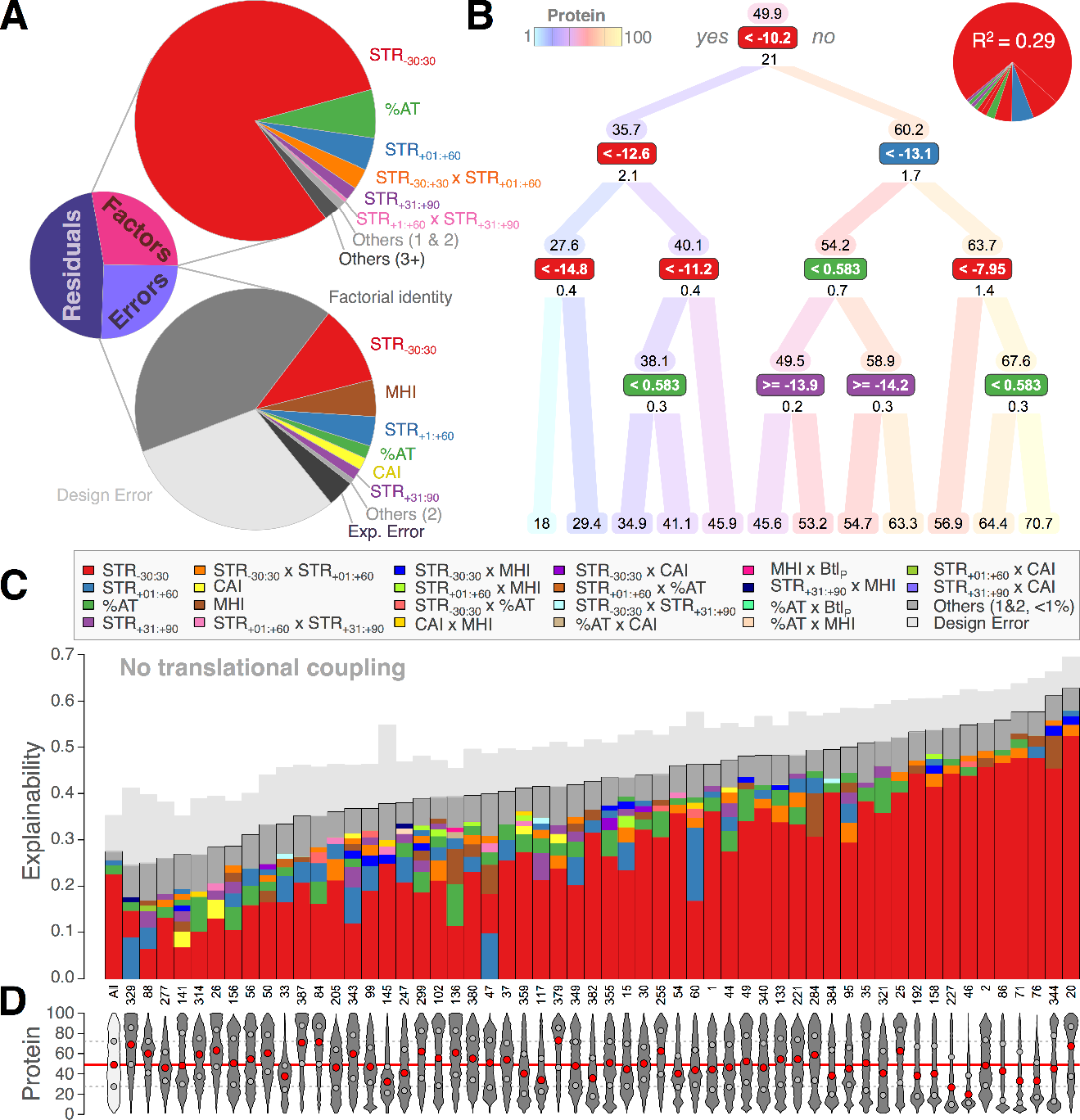
Secondary structures in the translation initiation region dominate protein production in the absence of coupling. **(A)** Decomposition of variance in P_NC_ across the whole library. The upper pie details the variance explained by the design properties and their interactions. The lower pie shows the contribution of various types of errors: experimental, within-series differences between close replicates (Design error), between-series differences (Factorial identity) and their second-order interactions with design properties (colored slices). Small effects are grouped in grey, distinguishing interaction depth (in parenthesis). **(B)** Recursive regression tree. At each node, data are split according to the rule shown in the colored box, which is heuristically chosen to maximize the explained variance. The property concerned with the rule is color-coded as in panels AC. R^2^ are shown below boxes and summarized in the upper-right pie. Average protein productions within each branch are shown above boxes and color-coded according to the upper-left thermometer. **(C)** Series-wise decomposition of explainable variance. For each series, effects >1% are stacked by decreasing values (colors as shown) and smaller effects are grouped on top (dark grey). The design error is shown above each bar (light grey). The explainable variance is the total variance subtracted by the experimental error. **(D)** Protein production profiles. Red and grey dots mark the median and interquantile range, respectively.

The dominance of secondary structures across the start codon reflects their negative impact on translation initiation (de Smit and van Duin, 1990; Kudla et al., 2009), which is generally regarded as the rate-limiting step in the translation process (Andersson and Kurland, 1990; Bulmer, 1991; Ciandrini et al., 2013; Shah et al., 2013). It follows that the effect of potential elongation-limiting properties, such as CAI, can only be observed in a context of high initiation (Supek and Šmuc, 2010; Tuller et al., 2010a; Figure S3G). Accordingly, we observe a strong negative correlation amongst factorial series between the contribution of initiation-limiting STR_−30:+30_ and that of all other properties (r=−0.72, Figure S3I).

### Epigenetic modulation of initiation rates via translational coupling reveals elongation determinants

We recently developed a standard bicistronic architecture that uses translational coupling from a defined leader cistron to mitigate the impact of mRNA structure on translation initiation of the downstream gene of interest (Mutalik et al., 2013a). Our reporter system is equipped with an improved version of the device that enables fully controllable disruption of the mRNA structure in the initiation region (Figure 1C and S4ABC). We performed the same experiment as above under condition of maximal coupling and were able to derive P_C_ for 238,458 strains (97,2% of the design library; Figure S4DE). Coupling leads to a drastic compression of protein production profiles toward the high end of the range observed previously (Figure 4D). Thus, for most variants in our library, the inducible coupling device enable effective increase in translation rates—without incurring any genetic alteration to the underlying sequences.

**Figure 4.**
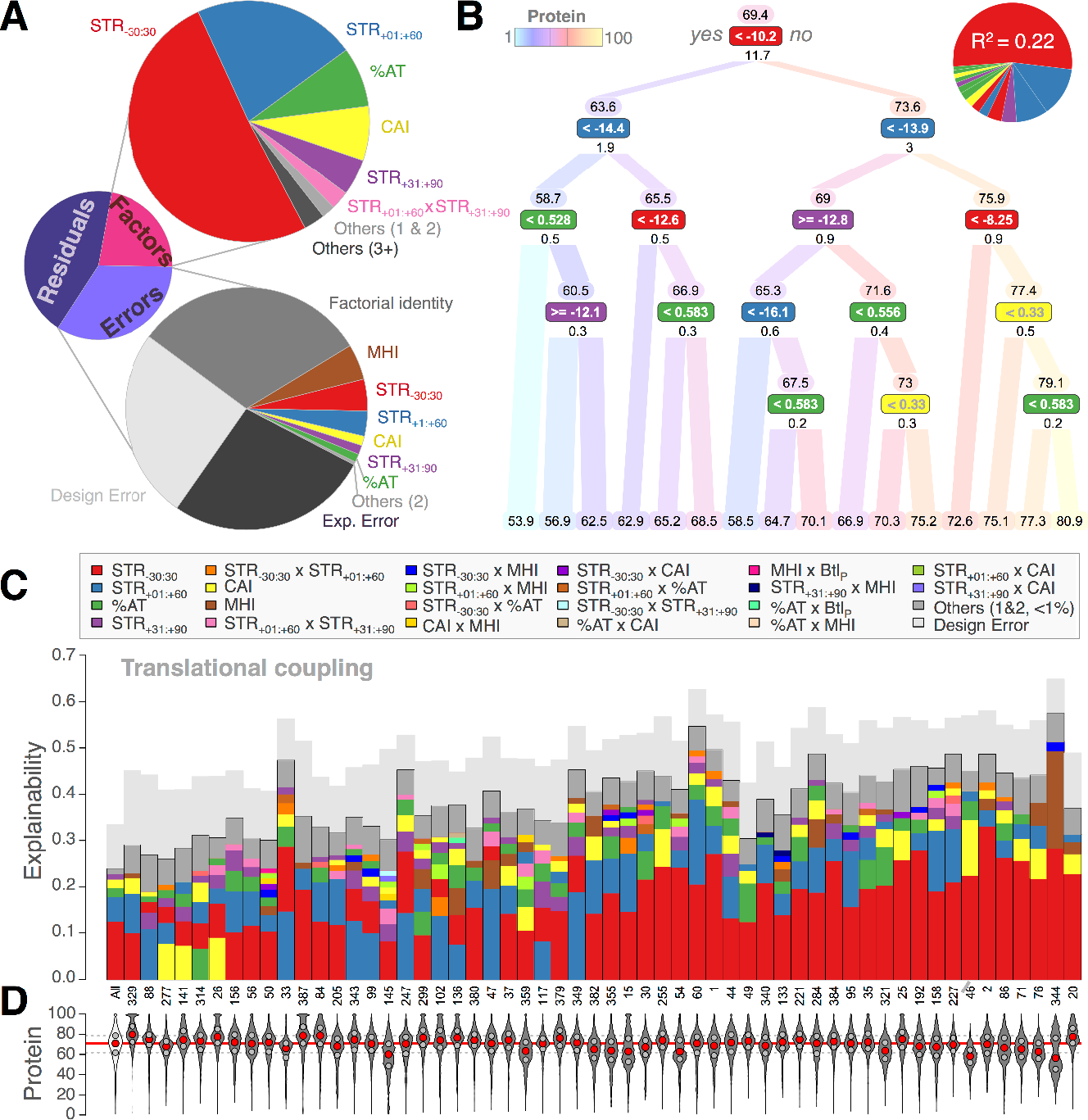
Translational coupling modifies the hierarchy of sequence properties impacting protein production. Same analysis as Figure 3 for P_C_. See Figure 3’s legend for details.

As previously, we conducted ANOVAs to decompose the contribution of design properties and various sources of error to P_C_ (Figure 4AC and S4FGH). Consistent with the reduced dynamic range of the response, the experimental error is more pronounced than before (9%; median CoV=0.06, Figure S4F). The design error is of the same magnitude (9%; median CoV=0.08) and the contributions of factorial identities and their interactions comparable to those observed for P_NC_ (11% and 6%, respectively). Altogether, the design factors and their second-order interactions explain 26-58% of the explainable variance within factorial series (39% on average, Figure 4D).

In spite of the coupling, structural properties persisted in showing dominant effects. The average contribution of STR_−30:+30_ across series nonetheless decreases from 60% to 38%. That of STR_+01:+60_, in contrast, rises from 7 to 18%. The farthest structure in the designed region (STR_+31:+90_) shows more modest improvements but exhibits substantial interactions with STR_+01:+60_ (5 and 2%, respectively). The fractions of explained variance contributed by %AT and MHI are similar to those observed in non-coupling conditions (7 and 4%). The contribution of CAI increases largely from 1 to 7%. The recursive regression procedure permits to further elucidate the hierarchical dependencies of these properties (Figure 4B). For example, the effect of CAI is restricted to sequences with weak STR_−30:+30_ and either weak STR_+1+60_ or medium STR_+1:+60_ associated with strong STR_+31+90_. Thus, initiation might still be limited by secondary structures in part of the library.

### Codon usage bias has a subdominant impact on elongation rate

P_C_ shows a strong and linear dependence on P_NC_ (Figure 5A), thereby providing an opportunity to single out properties that are specifically affected by the lift of initiation constraints. To that end, we conducted a multiple linear regression that incorporates P_NC_ as an explanatory variable in addition to the design properties (Figure 5B). In this framework, P_NC_ alone accounts for 61.8% of the variance in P_C_. STR_−30:+30_ and its interaction with P_NC_ represent insignificant parts of the residual variance (0.2% each). Hence, the unexpected impact of STR_−30:+30_ on P_C_ is a direct correlate of its contribution to P_NC_ and this structure does not seem to elicit a specific response under coupling. To a large extent, the same conclusion applies to %AT (0.2%). In sharp contrast, we find that STR_+01:+60_ and its interaction with P_NC_ account for substantial fractions of the residual variance (5.8% and 3.8%, respectively). CAI shows the next largest contribution (4.6%), while STR_+31:+90_ and its interactions with P_NC_ and STR_+1:+60_ also cumulate a sizeable effect (1.9% altogether).

**Figure 5.**
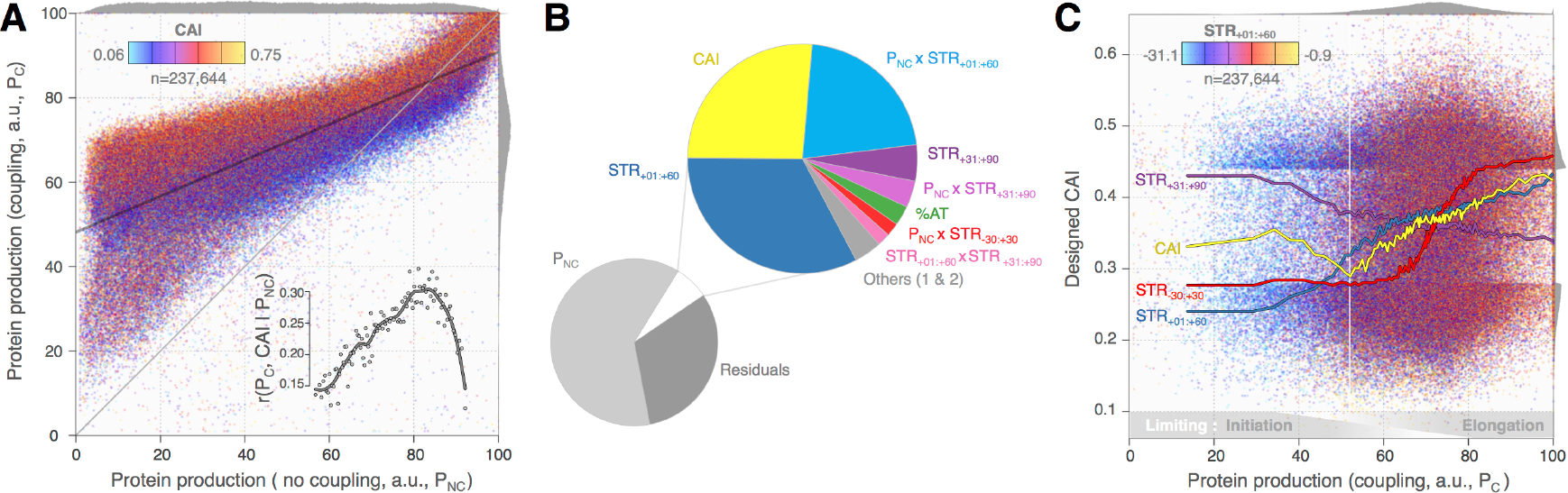
Translational coupling exposes elongation-limiting properties. **(A)** Impact of codon usage on elongation. Scatter plot of P_C_ versus P_NC_ colored by CAI, as shown. Distributions are shown on the edges. The linear regression of P_C_ on P_NC_ is plotted in dark grey. Inset shows rising correlations between P_C_ and CAI with increasing percentile of P_NC_. Higher codon adaptation supports commensurate increases in elongation rates when initiation is improved. **(B)** Decomposition of variance in P_C_ including P_NC_ as a predictor. The effects of design properties under normal translation are captured by P_NC_. Remaining contributions quantify differential effects elicited under coupling. Properties with effect <1% of the inset pie are grouped in grey. **(C)** Codon usage is subordinate to initiation. Scatter plot of CAI versus P_C_ colored by STR_+01:+60_, as shown. The median CAI for each percentile of P_C_ is plotted in yellow. Scaled equivalents for the median STR_−30:+30_ (red), STR_+01:+60_ (blue) and STR_+31:+90_ (purple) are shown for comparison. Past a production threshold (white line), CAI increases with P_C_ to support faster elongation. Below, initiation remains limited by strong STR_+01:+60._

For a given P_NC_ the increase in protein production that results from translational coupling is strongly correlated with CAI (Figure 5A). Thus, the lack of production improvement under coupling identifies sequences that are intrinsically limited by elongation. In other strains, increasing initiation reveals graded capacities to support heavier ribosomal fluxes that are commensurate to codon adaptation. The magnitude of this association first increases with P_NC_ (Figure 5A, inset), a trend that likely reflects the vanishing collateral impact of STR_−30:+30_ on elongation of the leader sequence (see next section). The association then sharply drops amongst highest protein producers, as *de facto* CAI distributions become increasingly skewed toward larger values (Figure S3G).

Excluding the bottom decile of protein producers, we find that P_C_ is associated with a smooth linear increase of the median CAI (Figure 5C). Amongst these strains, the median P_C_ increases substantially with CAI from 68 to 75 a.u. (r=0.16, Figure S5). That the lowest producers do not follow this association suggests that their translational regimes remains largely limited by initiation. As detailed below, this behavior is determined by strong STR_+1:+60_ combined with weak STR_+31:+90_. The distribution of CAI clearly follows these two structural properties (Figure 5C), in a pattern that reflects growing requirements for efficient elongation as initiation-limiting constraints are increasingly lifted.

### Versatile secondary structures dominate initiation and elongation rates

Relating the physical position of ribosomes at key steps of the translation process to the arrangement of the three overlapping designed structure windows (Figure 6A) permits to better grasp the diverse contributions of secondary structures to both P_NC_ and P_C_. Under coupling, ribosomes must unfold the entire STR_−30:+30_ and the first 16 nucleotides of the STR_+1:+60_. Re-initiation then entails accommodation of the SD_R_, as well as the precise adjustment of the reporter start codon onto the ribosomal P site (Milón and Rodnina, 2012; Yoo and RajBhandary, 2008). In our case, this involves a net backtracking of a single nucleotide. The STR_−30:+30_ being fully unfolded at this stage, its strong and seemingly indirect contribution to P_C_ is particularly surprising (Figure 4AB and 5AB).

Because potent STR_−30:+30_ mobilize the downstream half of the leader sequence (Figure 6A), such structures can be expected to slow elongation of the leader under coupling conditions. This would result in lower influxes of leader-bound ribosomes and subsequently lower coupling rates. Rather than a more direct but elusive mechanistic effect on reporter re-initiation, the impact of STR_−30:+30_ on P_C_ is thus likely to reflects the impact of secondary structures on leader elongation. Showing little contribution of CAI, P_NC_ is largely limited by initiation and constitutes an excellent proxy for the true strength of secondary structure in the initiation region. In this light, the prominent linear relationship between P_C_ and P_NC_ (Figure 5AB) suggests that secondary structures *in vivo* may have a stronger impact on elongation than codon-based features such as CAI.

The same reasoning is applicable to part of the STR_+01:+60_, since about half of an idealized fold centered in that window must be unfolded by a leader-bound ribosome (Figure 6A). However, the role of this structure in modulating initiation appears to be more direct. The long tails of low P_C_ observed in nearly all factorial series (Figure 4D) are characterized by strong STR_+01:+60_ (Figure 5C and 6C). Structures that nucleates after the footprint of leader-bound terminating ribosomes and are strong enough to extend toward the start codon may promote ribosomal backtracking, thereby hindering its precise positioning onto the start codon and eventually limiting re-initiation.

**Figure 6.**
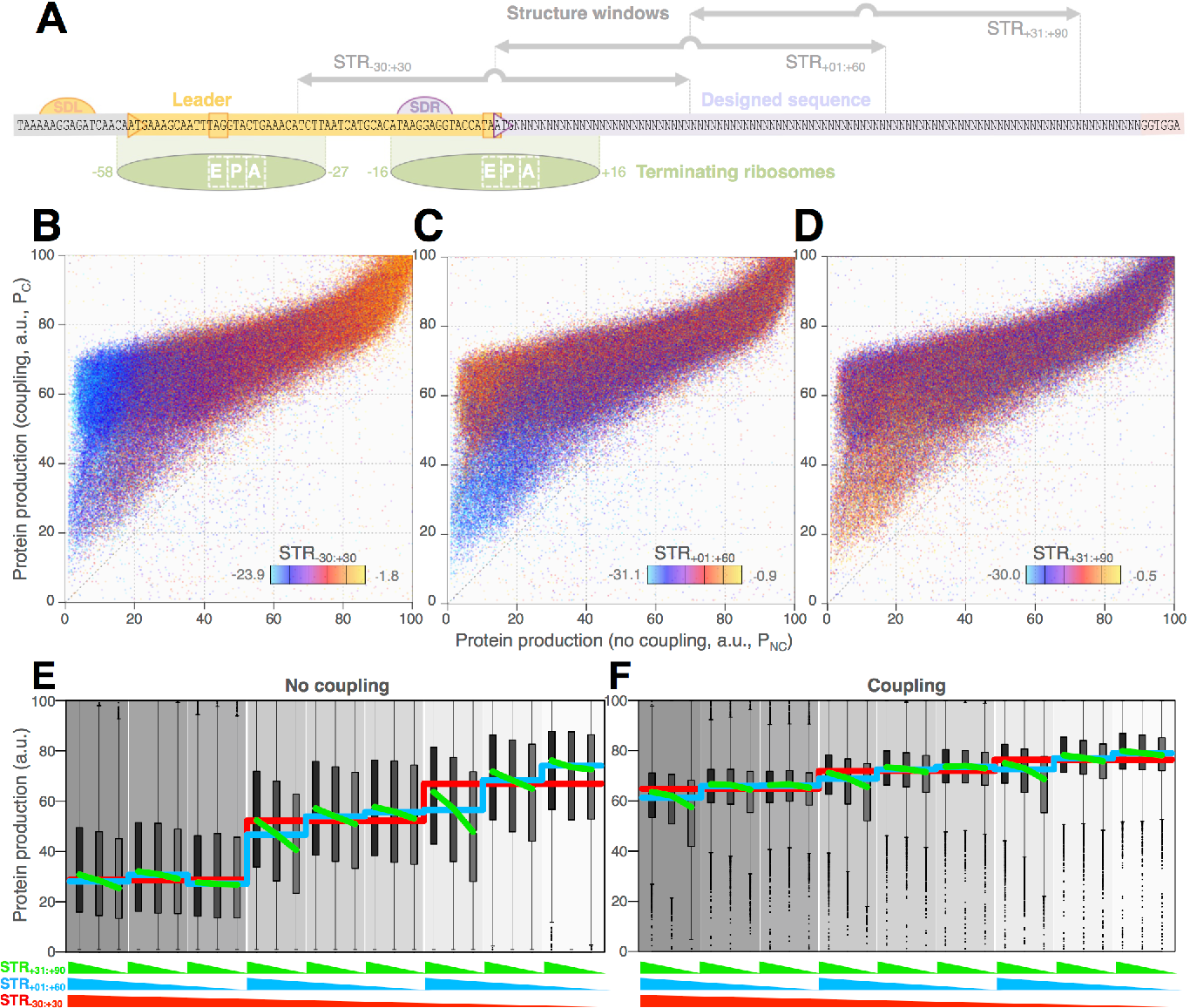
Complex interaction patterns between designed secondary structures. **(A)** Steric relationships between ribosomes and secondary structure. The organization of the reporter system and the position of design structures are detailed at scale. Green shapes show the precise footprint of leader-bound terminating ribosomes in non-coupling (left) and coupling (right) conditions. Positions are numbered from the reporter start codon. **(BCD)** Emergent co-distributions of structural properties in protein production spaces. Scatter plot of P_C_ versus P_NC_ colored by STR_−30:+30_ **(B)**, STR_+01:+60_ **(C)** and STR_+31:+90_ **(D)**, as shown. These plots highlight complex structural interaction patterns amongst low protein producers. **(EF)** Strong structural interactions impact protein production. These interaction plots outline the effect of all designed structural combinations on P_NC_ **(E)** and P_C_ **(F)**. Red lines mark the medians per level of STR_−30:+30._ Blue and green lines show the medians for combinations of STR_+01:+60_|STR_−30:%30_ and STR_%31:%90_|STR_−30:%30_|STR_+01:+60,_ respectively. Strong STR_+31:+90_ provide a dynamic switch capable of facilitating initiation through thermodynamic competition with STR_+01:+60._

The strong effect of STR_+01:+60_ requires further association with weak STR_+31:+90_ (Figure 6CD). In the mechanism outlined above, the destabilization of STR_+01:+60_ by terminating ribosomes releases part of the sequence involved in STR_+31:+90_. Strong STR_+31:+90_ that are capable of outcompeting the refolding of STR_+01:+60_ can therefore promote higher initiation by preventing the formation of this negative determinant. Such a thermodynamic competition may account for the otherwise puzzling positive association between STR_+31:+90_ strength and protein production (Figure 3B and 6EF), as well as for many interactions and hierarchical relationships identified above (Figure 3BC and 4BC). Likewise, one might expect strong STR_+01:+60_ to compete with initiation-limiting STR_−30:+30_ and therefore drive increased protein production. Our data do not evidence such antagonism, perhaps because such STR_+01:+60_ exert a sufficiently negative impact on initiation by themselves and substitute to the effect of the outcompeted STR_−30:+30_ (Figure 6EF). Accordingly, strains that undergo the most substantial protein production improvements under coupling are characterized by strong STR_−30:+30_ associated with weaker STR_+01:+60_ (Figure 6BC).

### Poor descriptions of secondary structures hinder accurate predictions

Given the dominant impact of structural properties on translation rates, we investigated whether we could define *post hoc* structural metrics to better describe our data. First, we computed minimum free energy predictions for windows of varied sizes positioned over the larger initiation region (−40:+81). This yielded smooth profiles of explanation, but only modest improvement over the original design windows (Figure 7A). Under coupling, this analysis corroborates the contribution of structures nucleating downstream of the pre-initiating ribosome footprint and capable of extending up to the start codon (Figure 7B). Again, we observed a substantial variability amongst factorial series (Figure 7CD), which mirrors the diversity of structure profiles associated with comparable protein productions (Figure S7). Altogether, these data suggest that a predictive approach based on a fixed window is not capable of capturing the diversity of folding situations.

To circumvent arbitrary window selection, we next computed predicted accessibilities for each nucleotide in the initiation region (−50:+45; Bernhart et al., 2006), and used them as so many predictors in a multiple regression analysis of protein production. This procedure affords better explanations of individual series, but is less accurate on the whole dataset (Figure 7I). While it does not provide any improvement from a predictive standpoint, this analysis clearly identifies pairing of the core SD_R_ motif and the ~25 nts after (but excluding) the start codon as the key determinants of translation initiation (Figure 7E). These data are in perfect agreement with functional genomics expectations(Figure 7F). The contribution of SD_R_ to P_C_ is markedly reduced (Figure 7GH), in agreement with the mechanism of translational coupling.

**Figure 7.**
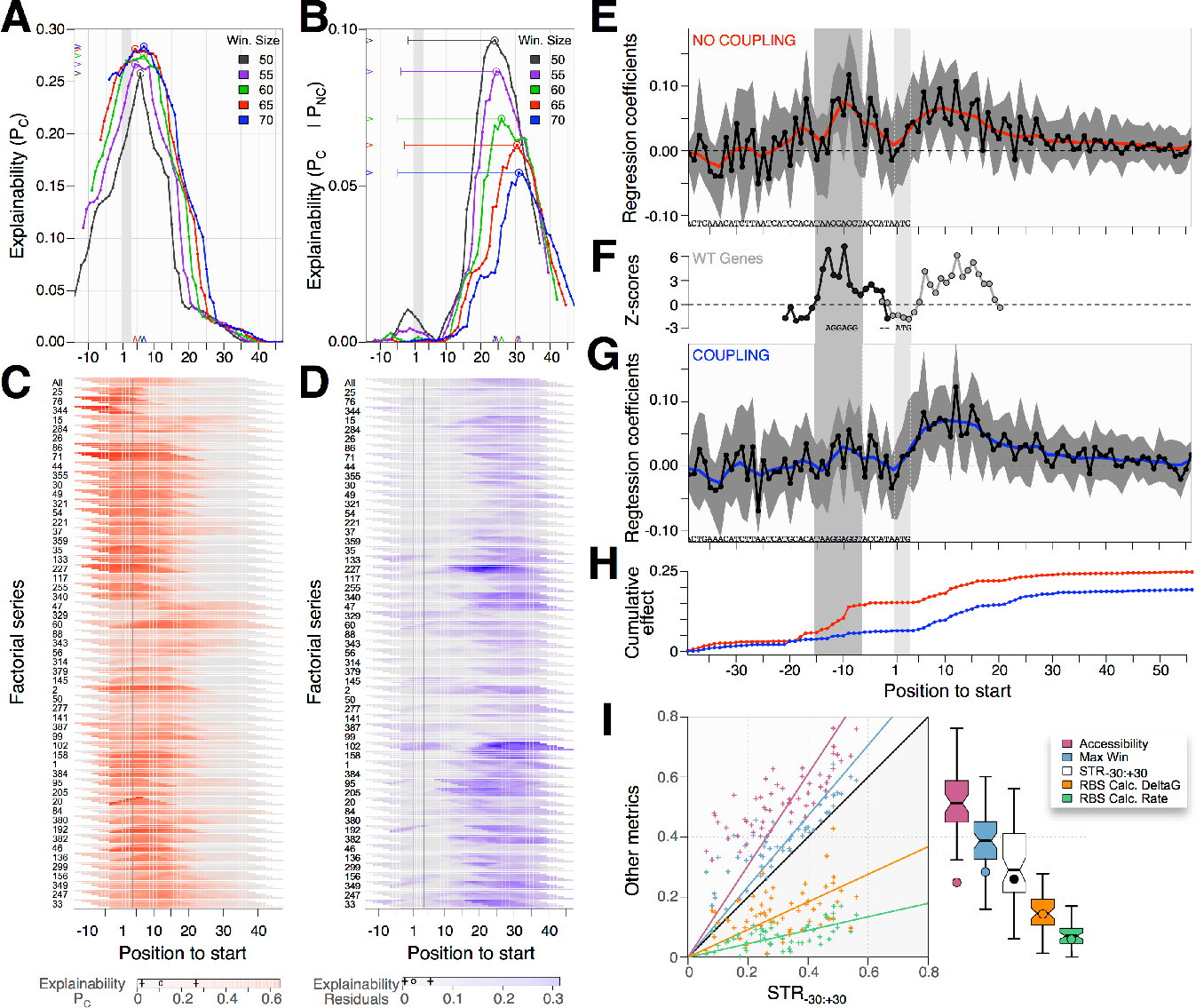
Lack of structural descriptors hinders predictions. **(A)** Modest explanatory improvements upon refinement of structural windows. Profiles obtained by regressing P_NC_ on MFE predictions for sliding windows of different sizes (color as shown). Circles and arrowheads mark profile optima. Longer windows are less sensitive to exact positioning. **(B)** Smaller windows at CDS start improve explanation under coupling. Profiles as in A but explaining residuals of the regression of P_c_ on P_NC_ (See Figure 5). Arrows mark halfwindow spans at profile optima. **(CD)** Highly variable profiles between series. Profiles in **A** and **B** are split by factorial series and shown as heatmap in **C** and **D**, respectively (thermometer as shown). Five rows within each series correspond to incrasing window length (top to bottom). **(E)** Functional impact of accessibility with nucleotide resolution. Shown are the medians (black) and interquartile ranges (grey) of standardized coefficients obtained upon regressing P_NC_ against average accessibility computed for every position within each series. The red line is a loess fit outlining the main trend. Dark and light grey background highlight SD_R_ and start codon, respectively. **(F)** Selection patterns toward increased accessibility mirror experimental results. Z-scores of mean accessibilities at each position computed for the WT *E. coli* genome with respect to shuffled genomes. **(G)** Translational coupling reduces the importance of SD_R_ accessibility. Same as E for P_C_. **(H)** Comparison of cumulative accessibility effects highlights lack of SD_R_ contribution to P_c_. **(I)** Predictions of translation output are limited. R^2^ from series-wise regressions of P_NC_ on STR_−30:+30_ are plotted against those obtained by regressing P_NC_ on nucleotide accessibilities (pink); best series-wise structure window (blue); composed minimum free energy (orange) andinitiation rate (emerald) from the RBS calculator (Espah Borujeni et al. 2014). Corresponding boxplots are shown. Circles mark R^2^ over the entire dataset.

## DISCUSSION

The ability to synthesize and assemble hundreds of thousands of precisely designed gene variants has allowed us to formally test a number of key theories about translational efficiency in *E. coli*.

We estimated that the properties implemented in our mDoE and their interactions could explain roughly a third of the measured variance in protein production. We could further link 6% of the unexplained variance to the inaccuracy of the property scores used for design. Significantly, we found that responses and explanation patterns are highly variable between factorial series. In fact, another 20% of global variance could be explained by indiscriminately considering the idiosyncratic behaviors that are specific to each series. In a few cases, it was possible to trace the sequence determinants of particular idiosyncrasies. For example, series 344 shows strongly bimodal distributions of protein production (Figure 3-4D). Many of its low producing strains comprise two contiguous prolines, an arrangement known to be translated very inefficiently (Chevance et al., 2014). The unusually high contribution of MHI in this particular series (Figure 3C and 4C) is then only incidental to the hydrophobic nature of these two residues (data not shown). This serve as a reminder that uncontrolled properties can confound even the most thorough mDoE.

The translation process is governed by the fine balance between the rates of initiation and elongation. The relative importance of these two steps has been a matter of active debate for decades, in part because most natural sequences have evolved to optimize their coordination. To disentangle these two processes, we have developed a new translational coupling device that enables controllable manipulation of initiation *in vivo*, without altering the sequences and their intrinsic elongation properties.

Some of the design properties have no measurable impact on protein production in either initiation or elongation-limiting conditions. In particular, our data do not lend support to the codon ramp hypothesis, which predicts that slowly translated codons at the beginning of genes should optimize the flow of ribosome and protein production (Navon and Pilpel, 2011). The MHI also shows negligible contributions. Yet, we investigated hydropathy because it produced one of the strongest associations with protein expression in our preliminary analysis of the *E. coli* genome (Figure S1DE). Further analyses suggest that our initial observation was confounded by functional constraints operating above regulatory level. Indeed, we found an overrepresentation of hydrophobic regions in the N-terminus of non-cytoplasmic proteins, which are overall less expressed than their cytoplasmic counterpart (data not shown). This confusion illustrates the fallibility of statistical inferences from natural sequences.

Properties describing secondary structures show prevalent contributions in all translation regimes. Strong structures capable of occluding the SD motif are critical initiation determinant and account for most of the variance in protein production. Secondary structures just downstream of the start codon can modulate initiation, especially when associated with weak, non-competing structures further downstream. Conversely, such competing downstream structures may assist unfolding and accommodation of the initiation region by the ribosome, hence unexpectedly increasing protein production. These results are strikingly in line with structural biases observed in the natural *E. coli* genome (Figure S1K). Such functional interactions thus appear significant enough to constrain the evolution of natural sequences. Finally, our data strongly suggest that transcript folding can strongly diminish elongation rates *in vivo*, as evidenced by the indirect impact on coupling efficiency of structure within the leader sequence.

Our results put forward the functional significance of complex intra-molecular folding transactions that operates dynamically as an mRNA is transcribed and simultaneously translated. Together with the diversity of structure contribution profiles between factorial series (Figure 7CD), they suggest that a good part of the design error and inability to fully explain protein production stems from limited capacities to adequately describe relevant structural features (Figure 7I). Further work will be needed in this respect to improve our predictive power.

We find that %AT contributes positively to protein production. These bases may favor the action of protein S1 of the 30S ribosomal subunit, which plays a key role in accommodating the transcript during initiation (Duval et al., 2013). Given that %AT remains slightly correlated to structural properties in our sequences (Figure 2E), it is also possible that this property captures a general structural feature that is not appropriately described by our measures of folding energy.

Finally, we show that improving initiation rates by translation coupling reveal a sizeable contribution of the CAI, which was largely hidden under normal conditions. The codon composition of a gene is expected to modulate its elongation rate due to the differential availability of cognate charged tRNA. That the effect of CAI is not detectable except in the highest protein producers strongly suggests that translation is natively limited by initiation in the vast majority of our library. Our results might contrast with the observation of natural sequences, the codon composition of which is probably evolutionary fine-tuned to match their respective initiation rates and thereby enable the production of necessary protein amounts. Only in this case does the CAI provide an accurate proxy for the global translation rate. It cannot currently be used as such with artificial sequences and its optimization is then subordinated to that of the initiation rate.

Altogether, our analyses provide important insight into the hierarchy of features that are important for translation efficiency. This permits a better understanding of the functional constraints that shape the evolution of natural sequences. Furthermore, it delineates a number of rules to optimize the translation of synthetic sequence for biotechnological applications. Given the poor predictive performances of our models and the underlying difficulties in properly capturing the impact of secondary structures, we refrained from developing an inaccurate predictive tool. More realistically, we propose a generic checklist procedure to maximize protein production that is largely inspired by the hierarchal classification of our data (Figure 3-4B): i) avoid any secondary structure capable of occluding the SD sequence; ii) avoid within-CDS structures capable of extending toward the start codon; iii) favor more distal structures capable of competing with their formation; iv) increase %AT in the first ~20 nucleotides of the gene; v) maximize codon usage in the rest of the gene, avoiding to introduce local structure that could slow elongation rate.

Protein production is perhaps the most immediate and relevant outcome of translation, but the process is deeply rooted in the global physiology of the cell. Fully understanding and controlling the repercussions of sequence perturbations therefore demands a more integrated and systemic analysis (Figure 1A) that is detailed ina companion paper.

## AUTHOR CONTRIBUTIONS

GC and APA conceived the work; GC and JCG designed the sequences; GC performed the experiment and processed the data; GC, JCG and APA analyzed the data; GC and APA wrote the manuscript.

## ACKNOWLEDGMENTS

We thank Jeff Sampson, Page Anderson and Steve Laderman from Agilent Laboratories for discussing OLS setup and processing; Chang Liu for providing the unnatural suppressor system; Vivek Mutalik, Laurent Jacob, Morgan Price, Michael Samoilov and Judy Savitskaya for useful discussions. We thank Agilent Laboratories and the Synthetic Biology Institute (SBI) for providing the OLS array. GC was funded by the Human Frontier Science Program (LT000873/2011-L); JCG by the Portuguese Fundacão para a Ciência e a Tecnologia (SFRH/BD/47819/2008); we acknowledge financial support by the Synthetic Biology Engineering Research Center (SynBERC). This work used the Vincent J. Coates Genomics Sequencing Laboratory at UC Berkeley (NIH S10 Instrumentation Grants S10RR029668 and S10RR027303).

## REFERENCES

Adhin, M.R., and van Duin, J. (1990). Scanning model for translational reinitiation in eubacteria. J. Mol. Biol. 213, 811–818.

Allert, M., Cox, J.C., and Hellinga, H.W. (2010). Multifactorial determinants of protein expression in prokaryotic open reading frames. J. Mol. Biol. 402, 905–918.

Andersson, S.G., and Kurland, C.G. (1990). Codon preferences in free-living microorganisms. Microbiol. Rev. 54, 198–210.

Bentele, K., Saffert, P., Rauscher, R., Ignatova, Z., and Blüthgen, N. (2013). Efficient translation initiation dictates codon usage at gene start. Mol Syst Biol 9, 675.

Bernhart, S.H., Hofacker, I.L., and Stadler, P.F. (2006). Local RNA base pairing probabilities in large sequences. Bioinformatics 22, 614–615.

Bulmer, M. (1991). The selection-mutation-drift theory of synonymous codon usage. Genetics 129, 897–907.

Cannarozzi, G.M., and Schneider, A. (2012). Codon Evolution (Oxford University Press).

Charneski, C.A., and Hurst, L.D. (2013). Positively Charged Residues Are the Major Determinants of Ribosomal Velocity. PLoS Biol 11, e1001508.

Charneski, C.A., and Hurst, L.D. (2014). Positive charge loading at protein termini is due to membrane protein topology, not a translational ramp. Molecular Biology and Evolution 31, 70–84.

Chevance, F.F.V., Le Guyon, S., and Hughes, K.T. (2014). The effects of codon context on in vivo translation speed. PLoS Genet. 10, e1004392.

Ciandrini, L., Stansfield, I., and Romano, M.C. (2013). Ribosome traffic on mRNAs maps to gene ontology: genome-wide quantification of translation initiation rates and polysome size regulation. PLoS Comp Biol 9, e1002866.

de Smit, M.H., and van Duin, J. (1990). Secondary structure of the ribosome binding site determines translational efficiency: a quantitative analysis. Proceedings of the National Academy of Sciences 87, 7668–7672.

Duval, M., Korepanov, A., Fuchsbauer, O., Fechter, P., Haller, A., Fabbretti, A., Choulier, L., Micura, R., Klaholz, B.P., Romby, P., et al. (2013). Escherichia coli ribosomal protein S1 unfolds structured mRNAs onto the ribosome for active translation initiation. PLoS Biol 11, e1001731.

Espah Borujeni, A., Channarasappa, A.S., and Salis, H.M. (2014). Translation rate is controlled by coupled trade-offs between site accessibility, selective RNA unfolding and sliding at upstream standby sites. Nucleic Acids Res. 42, 2646–2659.

Eyre-Walker, A., and Bulmer, M. (1993). Reduced synonymous substitution rate at the start of enterobacterial genes. Nucleic Acids Res. 21, 45994603.

Gingold, H., and Pilpel, Y. (2011). Determinants of translation efficiency and accuracy. Mol Syst Biol 7, 481.

Goodman, D.B., Church, G.M., and Kosuri, S. (2013). Causes and Effects of N-Terminal Codon Bias in Bacterial Genes. Science.

Grantham, R., Gautier, C., Gouy, M., Mercier, R., and Pavé, A. (1980). Codon catalog usage and the genome hypothesis. Nucleic Acids Res. 8, r49–r62.

Gu, W., Zhou, T., and Wilke, C.O. (2010). A universal trend of reduced mRNA stability near the translation-initiation site in prokaryotes and eukaryotes. PLoS Comp Biol 6, e1000664.

Guimaraes, J.C., Rocha, M., Arkin, A.P., and Cambray, G. (2014). D-Tailor: automated analysis and design of DNA sequences. Bioinformatics.

Gustafsson, C., Govindarajan, S., and Minshull, J. (2004). Codon bias and heterologous protein expression. Trends Biotechnol. 22, 346–353.

Ikemura, T. (1985). Codon usage and tRNA content in unicellular and multicellular organisms. Molecular Biology and Evolution 2, 13–34.

Ingolia, N.T., Weissman, J.S., Ghaemmaghami, S., Newman, J.R., Newman, J.R.S., and Weissman, J.S. (2009). Genome-Wide Analysis in Vivo of Translation with Nucleotide Resolution Using Ribosome Profiling. Science 324, 218–223.

Kapust, R.B., and Waugh, D.S. (2000). Controlled Intracellular Processing of Fusion Proteins by TEV Protease. Protein Expression and Purification 19, 312318.

Kosuri, S., Goodman, D.B., Cambray, G., Mutalik, V.K., Gao, Y., Arkin, A.P., Endy, D., and Church, G.M. (2013). Composability of regulatory sequences controlling transcription and translation in Escherichia coli. Proceedings of the National Academy of Sciences 110, 14024–14029.

Kudla, G., Murray, A.W., Tollervey, D., and Plotkin, J.B. (2009). Coding-Sequence Determinants of Gene Expression in Escherichia coli. Science 324, 255–258.

Li, G.-W., Burkhardt, D., Gross, C., and Weissman, J.S. (2014). Quantifying Absolute Protein Synthesis Rates Reveals Principles Underlying Allocation of Cellular Resources. Cell 157, 624–635.

Li, G.-W., Oh, E., and Weissman, J.S. (2012). The anti-Shine-Dalgarno sequence drives translational pausing and codon choice in bacteria. Nature 484, 538–541.

Milón, P., and Rodnina, M.V. (2012). Kinetic control of translation initiation in bacteria. Critical Reviews in Biochemistry and Molecular Biology 47, 334–348.

Mutalik, V.K., Guimaraes, J.C., Cambray, G., Lam, C., Christoffersen, M.J., Mai, Q.-A., Tran, A.B., Paull, M., Keasling, J.D., Arkin, A.P., et al. (2013a). Precise and reliable gene expression via standard transcription and translation initiation elements. Nat. Methods.

Mutalik, V.K., Guimaraes, J.C., Cambray, G., Mai, Q.-A., Christoffersen, M.J., Martin, L., Yu, A., Lam, C., Rodriguez, C., Bennett, G., et al. (2013b). Quantitative estimation of activity and quality for collections of functional genetic elements. Nat. Methods 10, 347–353.

Navon, S., and Pilpel, Y. (2011). The role of codon selection in regulation of translation efficiency deduced from synthetic libraries. Genome Biol 12, –.

Plotkin, J.B., and Kudla, G. (2010). Synonymous but not the same: the causes and consequences of codon bias. Nat Rev Genet 12, 32–42.

Pop, C., Rouskin, S., Ingolia, N.T., Han, L., Phizicky, E.M., Weissman, J.S., and Koller, D. (2014). Causal signals between codon bias, mRNA structure, and the efficiency of translation and elongation. Mol Syst Biol 10, 770–770.

Scott, M., Gunderson, C.W., Mateescu, E.M., Zhang, Z., and Hwa, T. (2010). Interdependence of cell growth and gene expression: origins and consequences. Science 330, 1099–1102.

Shah, P., Ding, Y., Niemczyk, M., Kudla, G., and Plotkin, J.B. (2013). Rate-Limiting Steps in Yeast Protein Translation. Cell 153, 1589–1601.

Sharon, E., Kalma, Y., Sharp, A., Raveh-Sadka, T., Levo, M., Zeevi, D., Keren, L., Yakhini, Z., Weinberger, A., and Segal, E. (2012). Inferring gene regulatory logic from high-throughput measurements of thousands of systematically designed promoters. Nat. Biotechnol. 30, 521–530.

Sharp, P.M., and Li, W.H. (1987). The codon Adaptation Index--a measure of directional synonymous codon usage bias, and its potential applications. Nucleic Acids Res. 15, 1281–1295.

Smanski, M.J., Bhatia, S., Zhao, D., Park, Y., B A Woodruff, L., Giannoukos, G., Ciulla, D., Busby, M., Calderon, J., Nicol, R., et al. (2014). Functional optimization of gene clusters by combinatorial design and assembly. Nat. Biotechnol. 32, 1241–1249.

Supek, F., and Šmuc, T. (2010). On relevance of codon usage to expression of synthetic and natural genes in Escherichia coli. Genetics 185, 1129–1134.

Taniguchi, Y., Choi, P.J., Li, G.-W., Chen, H., Babu, M., Hearn, J., Emili, A., and Xie, X.S. (2010). Quantifying E. coli proteome and transcriptome with single-molecule sensitivity in single cells. Science 329, 533–538.

Tuller, T., Waldman, Y.Y., Kupiec, M., and Ruppin, E. (2010a). Translation efficiency is determined by both codon bias and folding energy. Proceedings of the National Academy of Sciences 107, 3645–3650.

Tuller, T., and Zur, H. (2015). Multiple roles of the coding sequence 5' end in gene expression regulation. Nucleic Acids Res. 43, 13–28.

Tuller, T., Carmi, A., Vestsigian, K., Navon, S., Dorfan, Y., Zaborske, J., Pan, T., Dahan, O., Furman, I., and Pilpel, Y. (2010b). An Evolutionarily Conserved Mechanism for Controlling the Efficiency of Protein Translation. Cell 141, 344–354.

Tuller, T., Veksler-Lublinsky, I., Gazit, N., Kupiec, M., Ruppin, E., and Ziv-Ukelson, M. (2011). Composite effects of gene determinants on the translation speed and density of ribosomes. Genome Biol 12, R110.

Vogel, C., and Marcotte, E.M. (2012). Insights into the regulation of protein abundance from proteomic and transcriptomic analyses. Nat Rev Genet.

Welch, M., Govindarajan, S., Ness, J.E., Villalobos, A., Gurney, A., Minshull, J., and Gustafsson, C. (2009). Design parameters to control synthetic gene expression in Escherichia coli. PLoS ONE 4, e7002.

Yoo, J.-H., and RajBhandary, U.L. (2008). Requirements for translation re-initiation in Escherichia coli: roles of initiator tRNA and initiation factors IF2 and IF3. Mol. Microbiol. 67, 1012–1026.

